# Measuring capture, internalization and cytosolic delivery of extracellular vesicle-embedded syntenin

**DOI:** 10.64898/2026.02.05.704011

**Authors:** Lukas Hyka, Eliane Jaafar, Sofie Meeussen, Alain Joliot, Guido David, Pascale Zimmermann

**Affiliations:** Laboratory for Extracellular Vesicle Research, Department of Human Genetics, Katholieke Universiteit Leuven, Leuven, Belgium; Centre de Recherche en Cancérologie de Marseille (CRCM), Equipe Labellisée Ligue 2018, CNRS, Inserm, Institut Paoli Calmettes, Aix-Marseille Université, Marseille, France; INSERM U932, Institut Curie Centre de Recherche, PSL Research University, Paris, France

**Keywords:** EV capture, EV internalization, EV cargo delivery, quantitative assays, syntenin

## Abstract

Extracellular vesicles (EVs) mediate cell-to-cell communication and are considered potential drug delivery vehicles. Nevertheless, whether EV-embedded cargo can be efficiently delivered into the cytosol of recipient cells remains debated. Here, we investigated the fate of syntenin, a well-established internal cargo of small EVs (sEVs).

Using quantitative assays, we show that ∼85% of internalized sEV-embedded syntenin can be delivered to the cytosol of recipient cells within short periods of time. Yet, even at low dose, we find that the internalization of sEVs carrying syntenin is rather inefficient (∼0.03% of the administered dose). Moreover, we observe that the capture of sEVs by recipient cells is non-saturable over time and largely more efficient than their internalization. Finally, we identify the N-terminal domain of syntenin and the phosphorylation state of a Src-targeted tyrosine residue in this domain, as key determinants for its incorporation into sEVs that support cytosolic delivery.

These findings challenge, current views in the field by indicating that sEV internalization may be a marginal process (on the contrary to capture) and that cytosolic delivery can be highly efficient. Moreover, our study identifies molecular determinants governing cytosolic delivery of sEV-embedded syntenin.

## Introduction

Extracellular vesicles (EVs) are membrane-limited organelles released and taken up by all cell types. What remains a matter of debate is whether EV embedded or inner cargo can be efficiently delivered into the cytosol of EV recipient cells^1^.

Syntenin is a member of the PDZ domain-containing scaffold protein family that is involved in multiple cellular processes, including cell adhesion^2^, transcriptional activation^3^, and cell growth, proliferation, and cell cycle progression^4^. Importantly, using structure-function approaches, we established that syntenin is a key organizer of small extracellular vesicle (sEV) biogenesis, affecting up to 50% of the sEV population in the extracellular space^5–7^.

Structurally, syntenin consists of four domains: N-terminal domain, two central PDZ domains (PDZ1 and PDZ2), and a C-terminal domain, each mediating specific molecular interactions. In particular, the N-terminal domain of syntenin binds Alix via three repeats of the LYPXL motif (reminiscent of late viral domains implicated in viral budding). Together with Alix and syndecans, syntenin functions as a key component of sEV biogenesis by stimulating intraluminal vesicle accumulation in multivesicular bodies that support the secretion of sEVs^5^. Furthermore, the small GTPase ARF6 and its effector phospholipase D2 support this sEV biogenesis pathway, and the PDZ1 domain of syntenin binds the fusogenic lipid phosphatidic acid, the product of PLD2 activity^8^. Syndecans are abundant transmembrane proteins carrying heparan sulfate that strongly interact with the PDZ2 domain of syntenin and that support sEV biology in various manners^5,9,10^. In particular, we found that and syntenin support sEV-mediated pro-migratory activities upon phosphorylation of specific tyrosine residue by Src kinase in sEV producing cells^11^.

Despite the important role of syntenin in sEV biology and also its high abundance in these organelles ^5,12^, the fate of sEV-embedded syntenin, including its delivery into the cytosol of recipient cells, remains uninvestigated. Here, we use fluorescence and bioluminescence complementation assays to determine how efficiently sEV-embedded syntenin can be captured, internalized, and delivered into the cytosol of recipient cells. Moreover, we define molecular determinants that control cytosolic delivery of sEV-embedded syntenin.

## Materials and methods

### Reagents & antibodies

Fetal bovine serum (FBS, Sigma, 1681067), DMEM (Gibco, 41965039), XtremeGene 9 (Roche, 6365809001), Geneticin (Gibco, 10131027), EV-depleted FBS (FBS depleted of intrinsic EVs by overnight (18 h) centrifugation at 100,000 x g, followed by filtration through sterile 0.45 μm filters (Merck), Amicon 10 kDa filters (Merck, UFC901024), Furimazine (DC chemicals, DC70371). Cell mask deep red (ThermoFisher, C10046) staining was used according to the manufacturer’s instructions. The syntenin-1 antibody is homemade and was described before^2^. The antibodies against CD9, CD63 were kind gifts of Prof. Eric Rubinstein^13^ (Université Paris-Sud, Inserm UMRS 935, Villejuif, France). The anti-GFP antibody (Invitrogen, A11122) was used according to the manufacturer’s instructions. In house purified rabbit anti-LgBiT antibody obtained by immunization of full-length Nanoluciferase (NL) was provided by Alain Joliot and Clotilde Thery (INSERM U932, Institut Curie Centre de Recherche, PSL Research University, Paris, France).

Plasmids encoding NL and its split fragments (HiBiT and LgBiT) were kindly provided by Alain Joliot. The HiBiT fragment was fused to syntenin and CD63 by overlapping PCR and Gateway recombination cloning. GFP split fragments were kindly provided by Stephanie Cabantous (CRCT, Toulouse). All the fusions of different tags to syntenin and CD63 were N-terminal. All open reading frames were verified by sequencing.

### Cell culture and transfections

HEK293, and MCF-7 cells (ATCC) were cultured in DMEM supplemented with 10% FBS. For transfections, HEK293 cells were plated the day before so to achieve ∼40% confluency at the time of transfection. Cells were transfected with expression vectors using XtremeGene 9 according to the manufacturer’s protocol. Following transfection, cells were cultured for 48 hours, washed once with fresh culture medium (DMEM) before microscopy or secretome collections.

### Preparation of stable clones

Expression vectors were linearized. After 48 hours of transfection, MCF-7 cells were put under Geneticin selection at 500 μg/ml. These concentrations were determined as the minimal doses inducing full cell death after 14 days. Resistant colonies were picked after 21 days and expanded separately. Transgene expression in total cellular population was confirmed by confocal microscopy after immunostaining with anti-GFP or anti-LgBiT antibody.

### Preparation of EVs by differential ultracentrifugation (dUC)

For secretome collections cells were cultured in serum-free DMEM for 16 hours (5 ml medium per 10 cm plate of 1.2 million cells (plated on day 1)). Conditioned media were collected after 16 hours and subjected to three sequential centrifugation steps at 4°C: 10 min at 1,500 x g (50 ml Falcon tube, rotor Sigma 12310), to remove cells and cell debris; 30 min at 10,000 x g (10K; 50 ml Falcon tube, rotor Sigma 12310) to pellet large particles (lEVs); and 1 h at 100,000 x g (100K; Beckman Open-Top Thinwall Ultra-clear Tube rotor SW32Ti) to pellet small particles (sEVs). 100K pellets were washed in PBS, centrifugated for 1 h at 100,000 x g (Beckman Microfuge Tube Polyallomer, rotor TLA-55), and re-suspended in 40 μl of PBS, as were 10K pellets for further analysis by Western blot. The corresponding cell layers were kept on ice, scraped in lysis buffer (Tris-HCl 25 mM, pH 7.4, NaCl 150 mM, EDTA 1 mM, NP40 1% and protease inhibitors), and incubated for 45 min at 4°C under gentle agitation. Cell lysates were cleared by centrifugation at 10, 000 x g at 4°C for 10 min before analysis by Western blot.

### Preparation of EVs by concentrating conditioned medium (CCM sEVs)

For secretome collections, cells were cultured in serum-free DMEM for 16 hours (5 ml medium per 10 cm plate of 1.2 million cells (plated on day 1)). Conditioned media were collected and processed as for dUC, for cell debris and lEVs removal. The 10K pellet supernatant was loaded on Amicon 10 kDa filters and concentrated by centrifugation at 2,000 x g until the volume dropped from ∼10 ml to ∼300 μl (33 x concentration). This concentrated supernatant was referred to as the CCM sEV fraction and characterized by Western blot or used in cell incubation experiments. For the comparison of GFP11 sorting with dUC (Fig. 1c, Supplementary Fig. 2) conditioned media were pooled upon collection and then again separated. sEVs were then prepared as described in dUC or CCM sEVs preparation sections.

**Figure 1.**
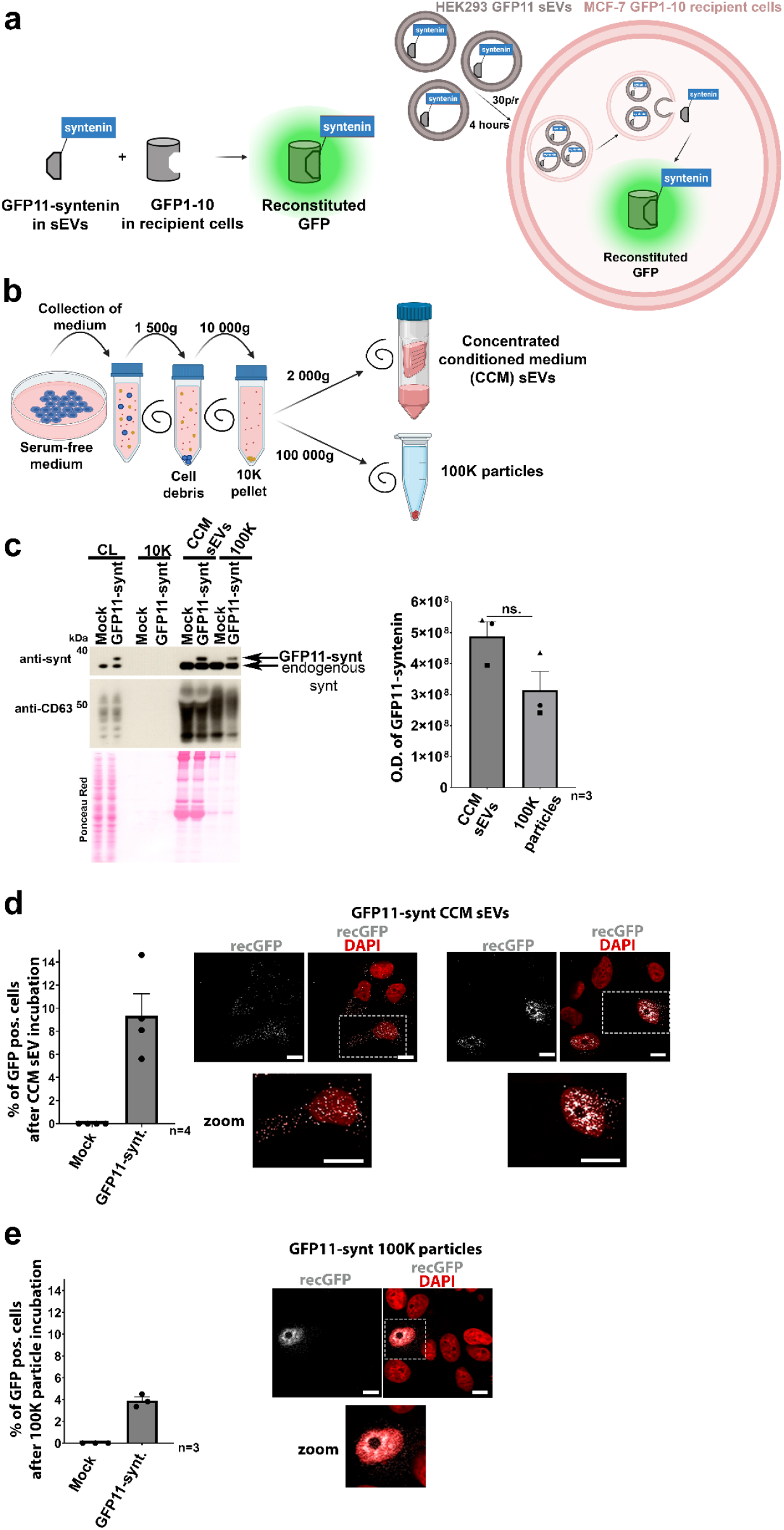
GFP11-syntenin is sorted to small extracellular vesicles (sEVs) and supports reconstitution of GFP in recipient cells expressing GFP1-10. **(a)** Schematic representation of the GFP complementation assay. GFP can be split into two non-fluorescent fragments (GFP11 and GFP1-10). In our setting, GFP1-10 is stably expressed in the cytosol of recipient MCF-7 cells, GFP11 is fused to syntenin and present in the sEVs of donor HEK293 cells. Fluorescence is reconstituted when the two fragments come in direct contact. MCF-7 cells stably expressing GFP1-10 were incubated with HEK293-derived sEVs at a producing:recipient cell ratio of 30:1 (30p/r). Cells were incubated for 4 hours with the 100K pellet or CCM sEVs of HEK293 cells transiently overexpressing GFP11-syntenin or Mock as control. **(b)** HEK293 sEVs fractions were prepared by differential ultracentrifugation or by concentrating the conditioned media (CCM sEVs) and analyzed by Western blot using anti-syntenin antibodies **(c, left)**. Note that the syntenin signals (endogenous or GFP11-fusion) are present in the 100K and CCM sEVs fractions but absent from 10K pellets. Cell lysates (CL) correspond to 20,000 cells, secretome fractions were prepared from 2.4 million cells. Endogenous syntenin and CD63 were used as markers and loading controls. **(c, right)** Bar graph showing average band intensities + SEM of GFP11-syntenin (Y-axis, integrated density) prepared by concentration (CCM sEVs) or by differential ultracentrifugation (100K particles), as indicated. Individual points correspond to individual western blot assays shown in Supplementary Fig. 3. **(d, e)** Left, bar graphs showing the average percentages + SEM of GFP positive MCF-7 GFP1-10 cells after the incubation with Mock or GFP11-syntenin CCM sEVs (d) or 100K particles (e). Individual data points correspond to independent biological repeats and represent values obtained from at least 40 nuclei scored at random. Right, confocal micrographs showing the localization of the GFP fluorescence (grey) in MCF-7 GFP1-10 cells treated with GFP11-syntenin CCM sEVs (d) or 100K particles (e). Nuclei were stained with DAPI (red). Inserts correspond to higher magnifications of indicated areas. Note dotty patterns in the cytosol (left) and the nuclei (right) of the reconstituted GFP followed by CCM sEV incubation **(d)** and nuclear reconstituted GFP patterns followed by 100K particle incubation **(e)**. Scale bar is 10 μm. Related to **Supplementary Figs. 1, 2 and 3**.

**Figure 2.**
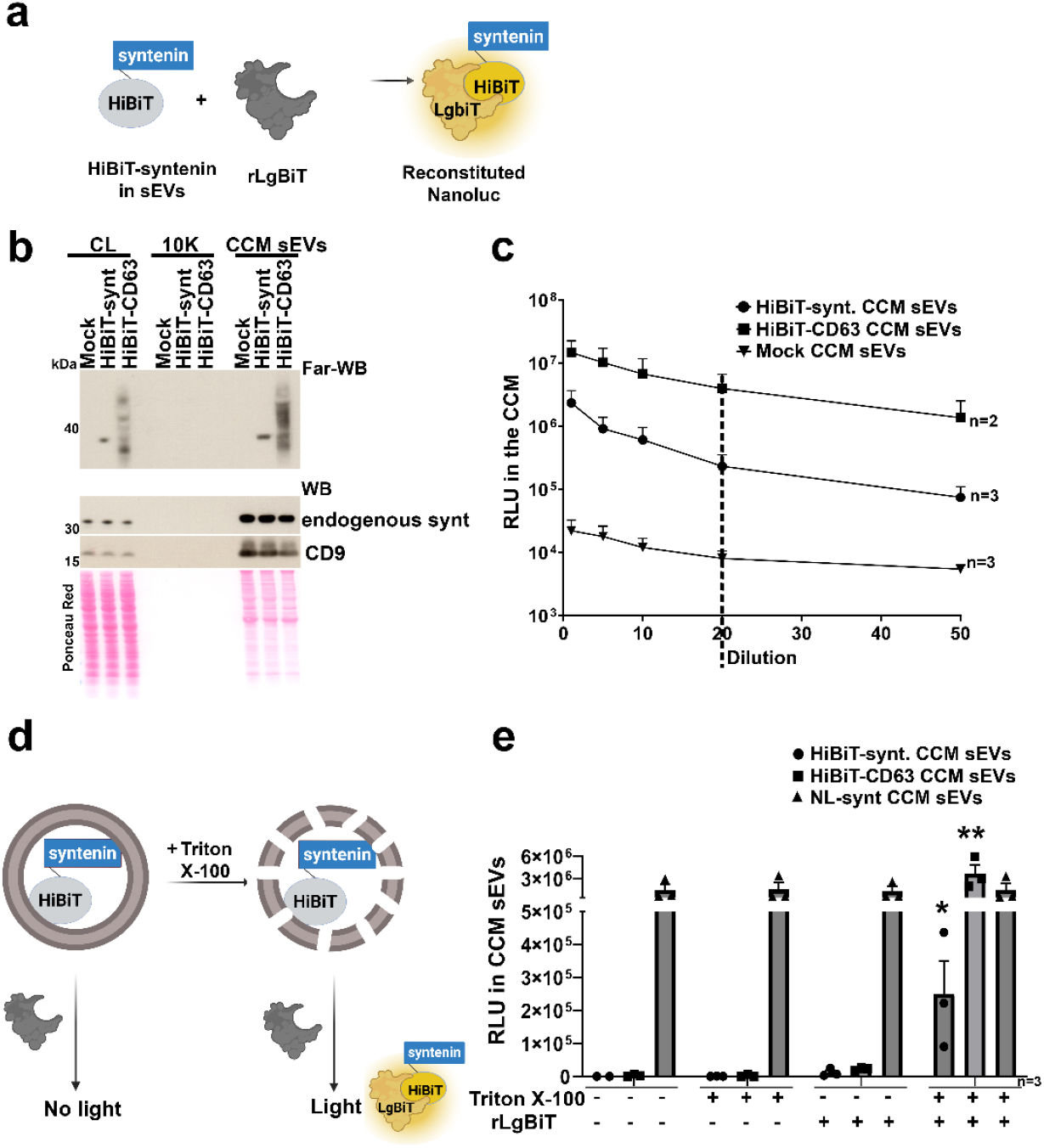
All HiBiT-syntenin present in the secretome of HEK293 cells is encapsulated within small extracellular vesicles. **(a)** Schematic representation of the Nanoluciferase (NL) reconstitution. NL can be split into two non-functional fragments (HiBiT and LgBiT) that reconstitute enzymatic activity upon direct interaction. In this figure, HiBiT is fused to syntenin (CD63, not shown). HEK293 cells are transfected with expression vectors for these constructs and their conditioned media are processed to prepare HiBiT-containing CCM sEVs. **(b)** Top part, Far Western blot analysis of cell lysates (CL) and secretome fractions (10K pellet, CCM sEVs) from HEK293 cells transiently expressing HiBiT-syntenin, HiBiT-CD63, or empty vector (Mock). Blots were probed with recombinant LgBiT (rLgBiT) before Furimazine substrate was added, to reconstitute NL and generate luminescence signals. CL corresponds to 20,000 cells, secretome fractions were prepared from 3.6 million cells. Bottom part, control Western blots illustrating the presence of endogenous syntenin and CD9 signals. Ponceau red staining served as a loading and transfer control. **(c)** Graph showing the average luminescence signals (RLU, Y-axis), + SEM (SE for CD63), obtained after various dilutions of CCM sEVs containing different constructs as indicated. Signals were obtained after incubation with Furimazine in the presence of 0.05% Triton X-100 and 2.5 μg rLgBiT. See Supplementary Figs. 3b and 3c for set-up assays. The dashed line indicates the dilution selected for total CCM sEV dose measurements in further experiments. The n-number indicates the number of independent biological repeats. **(d)** Schematic representation showing the experimental setup to verify the intraluminal location of HiBiT-tagged constructs (illustrated for syntenin). HiBiT and LgBiT can reconstitute into an enzymatically active form when not separated by a membrane. To determine the internal/external orientation of HiBiT-tagged syntenin, CCM sEVs were incubated with rLgBiT in the presence or absence of Triton X-100, followed by Furimazine substrate addition (not shown) and luminescence measurement. The n-number indicates the number of independent biological repeats. **(e)** Bar graph showing average luminescence signals (relative light units, RLU, Y-axis) + SEM obtained from CCM sEVs containing HiBiT-syntenin, HiBiT-CD63, or NL-syntenin (used as positive control). CCM sEVs were treated or not with Triton X-100 and/or rLgBiT, as indicated at the bottom. The membrane-permeant substrate, Furimazine was used throughout. Note that luminescence signal for HiBiT-fusions was detected only upon membrane permeabilization and rLgBiT addition, indicating intraluminal localization of the HiBiT tag. As expected, NL-syntenin luminescence signal was unaffected by the presence of Triton X-100 and/or rLgBiT. Each point represents an independent biological replicate. One-way ANOVA with Tukey’s correction for multiple comparisons (HiBiT data only) was applied. Related to **Supplementary Fig. 4**.

### Immunofluorescence analyses

The cells were cultured on eight-well chamber slides (Nalgene Nunc, 177402) overnight, fixed with 4% PFA for 15 min, washed with PBS, and permeabilized with 0.05% saponin in PBS for 20 min, in the presence of 0.3% BSA for blocking non-specific reactions. After incubation with primary antibodies, cells were incubated with Alexa-conjugated secondary antibodies (Invitrogen). Cells were visualized either by nuclei DAPI staining or by cell membrane staining using Cell mask deep red followed by mounting with ProLong Glass Antifade Mountant (ThermoFisher, P36982). Samples were observed with an Olympus Fluoview 1000 confocal microscope. Images were analyzed using ImageJ/Fiji (National Institute of Health, Bethesda, MD) and Photoshop (Adobe, San Jose, CA) software.

### (Far) Western Blotting

Cleared cell lysates and EV-enriched fractions were resuspended in PBS, boiled for 5 min in Laemmli buffer (10% glycerol, 2% SDS, 10 mM Tris, 10 mM EDTA, bromophenol blue) and fractionated by SDS-PAGE (NuPAGE 4-12% Bis-Tris gels, Invitrogen) before transfer to nitrocellulose membranes (Hybond-C extra, 0.45 μm, GE Healthcare). Cell lysates loaded always corresponded to approximately 20 000 plated cells (day 1) – equivalent to 20 μg protein according to bicinchoninic acid assay (BCA). EV-enriched fractions were collected from 2.4 million plated cells (day 1), unless specified differently. Membranes were probed with primary and HRP-conjugated secondary antibodies or incubated with 11.5 μg/ml recombinant LgBiT (rLgBiT) and 5 μM of Furimazine substrate as described in^14^, followed by chemiluminescence detection (Perkin Elmer). Band intensities were quantified using ImageJ by measuring integrated density after background subtraction.

### EV uptake (capture/internalization) experiments

#### GFP11-syntenin delivery

CCM sEVs prepared from HEK293 cells transiently overexpressing GFP11-fusions were incubated in serum-free DMEM for 4 hours with recipient MCF-7 GFP1-10 cells at a ratio of 30 producing cells per recipient cells (30p/r) as counted by Countess 3 automatic cell counter (ThermoFisher; 16812556) at the time of secretome collection (HEK293 cells) and at the time of sEV addition (MCF-7 cells). Recipient cells were plated on 8-well glass chamber slides the day before and washed with serum-free medium before the start of the experiment. Following the incubation with CCM sEVs or 100K particles, recipient cells were washed with PBS and prepared for confocal microcopy as described above. DAPI or Cell mask deep red signals were used to determine the GFP positivity percentage, all counted values can be found in Supplementary Table 1.

#### Full HiBiT dose measurement

To measure the total dose of HiBiT-sEV, an aliquot (1/20) of the CCM sEVs preparation was taken and incubated with 2.5 μg of rLgBiT + 0.05% of Triton X-100 followed by Furimazine substrate for 5 min as determined in ^14^ before the luminescence measurement.

#### HiBiT/NL fusion sEV delivery

CCM sEVs of NL- or HiBiT-fusion overexpressing HEK293 cells were incubated with MCF-7 LgBiT or wild-type recipient cells at 30p/r (see above), for different periods of time, as indicated. Cells were washed three times with versene (Gibco), trypsinized, unless specified differently, pelleted and washed three times with OptiMEM (Fisher sciences) at 500 x g for 5 mins at 4°C, followed by incubation with Furimazine for 5 min before the luminescence measurement on a Victor Nivo plate reader (PerkinElmer). For the cytosolic delivery, the cells were split in two halves and these halves were washed three times with OptiMEM. One half was used to measure the EV uptake/internalization by incubating the cells with 2.5 μg of rLgBiT in the presence of 2% of Triton X-100 after setup (see Supplementary Figs. 5c, d) and 8.3 μM of Furimazine for 5 min before measurement of luminescence. The other half was used to measure the cytosolic delivery by incubating the cells solely with membrane-permeant Furimazine before measurement of luminescence. For simplified cytosolic delivery assay cells were washed twice before trypsin treatment and once after trypsin treatment. For the measurement of the ‘cytosolic delivery’ fraction, cells were incubated solely with Furimazine. For the measurement of the ‘total sEV uptake’ fraction, cells were incubated with 0.8% of Triton X-100 followed by addition of Furimazine substrate. Each CCM sEV preparation served for 1-3 independent measurements of internalization and cytosolic delivery with at least 2 technical repeats.

### Statistical analysis

Statistical comparisons were performed using ANOVA followed by Tukey’s or Dunnet’s multiple comparison test or Student T-test. Graphs depict means or medians with SEM or maximal and minimal values from at least three independent experiments, unless otherwise stated. Statistical analyses were conducted using GraphPad Prism v9.0.

## Results

### sEVs fractions from HEK293 cells overexpressing GFP11-syntenin support GFP fluorescence reconstitution in MCF-7 cells expressing GFP1-10

To determine whether sEV-embedded syntenin is delivered to the cytosol of recipient cells, we first used a Green Fluorescent Protein (GFP) complementation assay. This assay works on the principle that GFP can be split into two non-fluorescent fragments, a 16 amino-acid peptide (GFP11) and a 214 amino-acid peptide (GFP1-10)^15^ (Fig. 1a, left). Cytosolic delivery of sEVs containing GFP11-syntenin into GFP1-10-expressing recipient cells is expected to result in GFP reconstitution, thereby providing a direct spatial readout for cytosolic delivery of syntenin (Fig. 1a, right). We first prepared MCF-7 cells stably expressing GFP1-10 (Supplementary Fig. 1a) and confirmed that upon transfection of expression vectors for GFP11 fusion constructs, we detect GFP fluorescence in these cells (Supplementary Fig. 1b). Transfection of an expression vector encoding GFP11 peptide alone (Supplementary Fig. 1c) did not lead to GFP fluorescence, consistent with the reported instability of this peptide when taken in isolation^16^.

Having validated the assay, we transfected HEK293 cells (sEV donor cells) with either an empty vector (Mock), an expression vector encoding GFP11-syntenin as the sEV cargo of interest, or an expression vector encoding GFP11-CD63, which has been previously reported to undergo cytosolic delivery and was therefore included as a reference cargo for comparison^17,18^. HEK293 conditioned media were cleared for cell debris and large EVs (lEVs, 10K particles) and either concentrated by low-speed centrifugation on columns or ultracentrifugation. The concentrated conditioned medium is further referred to as CCM sEVs and the ultracentrifuged pellet is further referred to as 100K particles (Fig. 1b). Expression of the different constructs, and their specific sorting in CCM sEVs or 100K particles was analyzed by Western blot (Fig. 1c, Supplementary Fig. 2). While no signal was observed in the 10K pellet, GFP11-syntenin was found to accumulate in CCM sEVs, and in 100K particles.

We then incubated HEK293-derived CCM sEVs or 100K particles (derived from the same conditioned medium) with MCF cells expressing GFP1-10, at a dose of 30 producing (p) HEK293 cells per recipient (r) MCF7 cell (30p/r) for 4 hours (Fig. 1a) and assessed the reconstitution of GFP fluorescence. As expected, we observed no GFP signal with the Mock (Fig. 1d, e, Supplementary Fig. 3a), and GFP11-CD63 CCM sEVs and 100K particles supported GFP reconstitution (Supplementary Fig. 3b, c). GFP reconstitution was also readily detected upon incubation with GFP11-syntenin CCM sEVs (Fig. 1d, e). Quantification revealed that 9.4 +/-1.9% of MCF-7 GFP1-10 cells incubated with GFP11-syntenin CCM sEVs displayed GFP fluorescence (Fig. 1d, left), whereas only 3.9 +/-0.3% of the cells incubated with GFP11-syntenin 100K particles were GFP positive (Fig. 1e, left). Notably, GFP signals induced by GFP11-syntenin CCM sEVs appeared concentrated in dotty intracellular patterns, whereas GFP signals induced by GFP11-syntenin 100K particles had dispersed nuclear localization (Fig. 1d, e).

**Figure 3.**
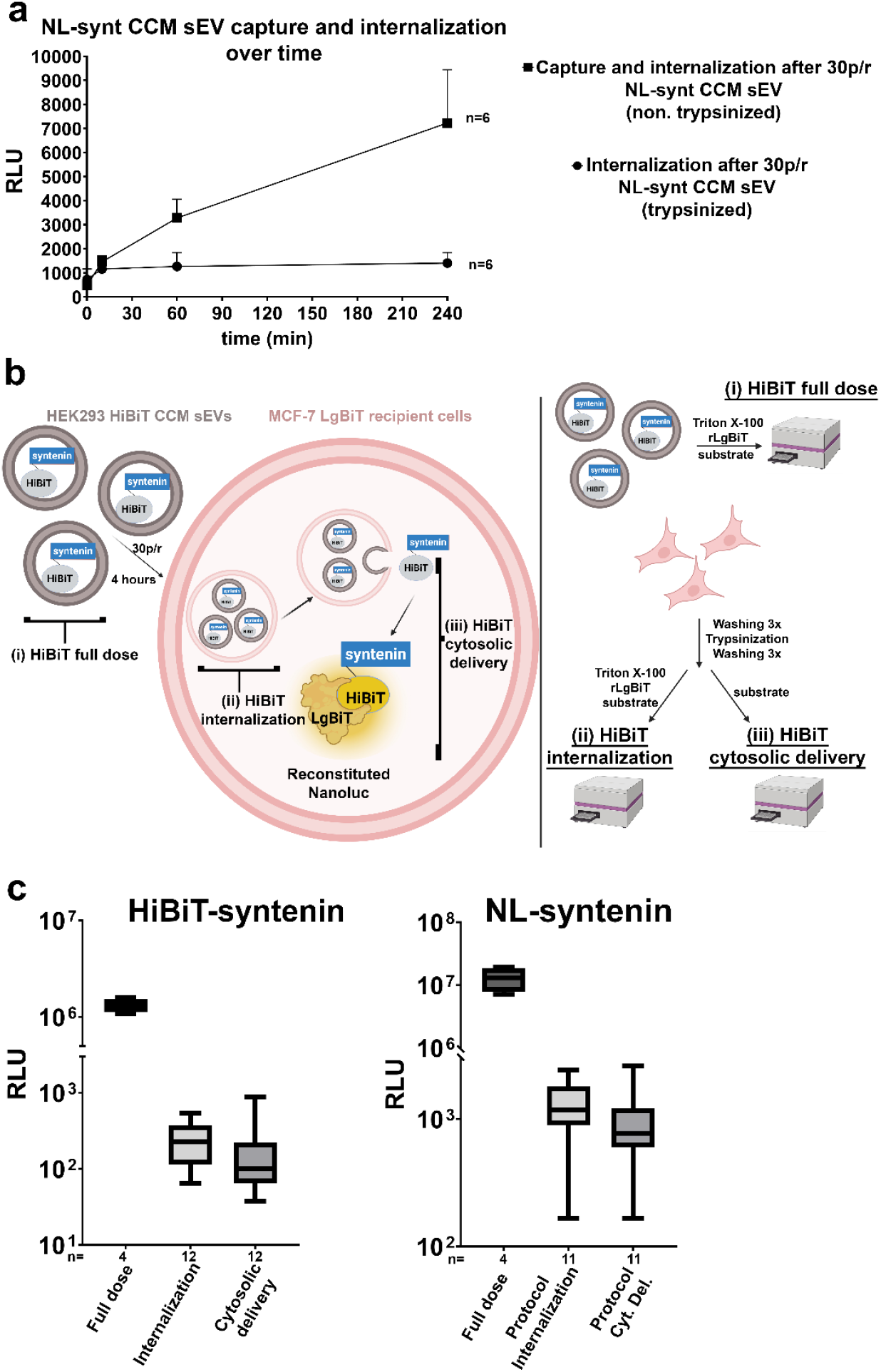
Capture versus internalization of sEV NL-syntenin and measurements of sEV HiBiT-syntenin full dose, internalization and delivery to the cytosol of recipient cells by NL complementation assay. **(a)** Average luminescence signals (relative light units, RLU, Y-axis) + SEM obtained from MCF-7 WT cells after incubation at 37°C, for different periods of time, with CCM sEVs (30p/r) containing NL-syntenin. After incubation, cells were trypsinized or not, as indicated. Signals measured after washing (in the absence of trypsin) were taken as a measure of sEV capture together with internalization, whereas signals measured after trypsinization and washing were taken as a measure of sEV internalization only. The luminescence signals were measured after addition of the membrane-permeant substrate Furimazine. Note, that capture increases continuously over time, whereas the internalization fraction rapidly reaches a plateau value. The n-number indicates the number of independent biological repeats. **(b)** Schematic overview recapitulating the different measurements (left) and the experimental approach (right). Briefly, for the determination of the full dose (i), an aliquot of the CCM sEVs was incubated with rLgBiT in the presence of Triton X-100 before Furimazine addition. For the determination of the internalization (ii) and the cytosolic delivery (iii), MCF-7 LgBiT were incubated with CCM sEVs (30p/r) for 4 hours, washed and trypsinized. Samples were split in two equal fractions. The first fraction was incubated with Triton X-100, rLgBiT before Furimazine addition to measure the internalized sEVs. The second fraction was solely incubated with Furimazine to measure the sEV cytosolic delivery. **(c)** Bars and whiskers graphs showing the median luminescence signals with minimal and maximal values, obtained with CCM sEVs containing HiBiT-syntenin (left) or NL-syntenin used as positive control (right). Background signals from MCF-7 LgBiT cells treated with Mock CCM sEVs were subtracted. Each CCM sEV preparation served for 2-3 independent measurements of internalization and cytosolic delivery with at least 2 technical repeats. Related to **Supplementary Fig. 5**.

Taken together, these data indicate that sEV syntenin can be delivered to the cytosol of recipient cells, and that ultracentrifugation may alter this delivery.

### Nanoluciferase complementation assays measuring CCM sEV HiBiT-syntenin dose

To complement these GFP assays with a more quantitative approach, we used Nanoluciferase (NL) based assays^19,20^. Similar to GFP, NL can be split into two non-functional fragments: HiBiT (11 amino acids) and LgBiT (159 amino acids) (Fig. 2a). We prepared expression vectors encoding HiBiT fused to the N-terminus of syntenin or CD63 as a reference cargo. Cell lysates (CL), lEVs (10K), and CCM sEVs from HEK293 cells transiently overexpressing these constructs were analyzed for the presence of HiBiT-tagged proteins by Far-Western blot using recombinant LgBiT (rLgBiT) and Furimazine as substrate (Fig. 2b, Supplementary Fig. 4a). Signals at the expected molecular weights were detected in CL and CCM sEVs but not in lEVs (10K), indicating selective enrichment of HiBiT-tagged syntenin and CD63 in sEV fractions.

**Figure 4.**
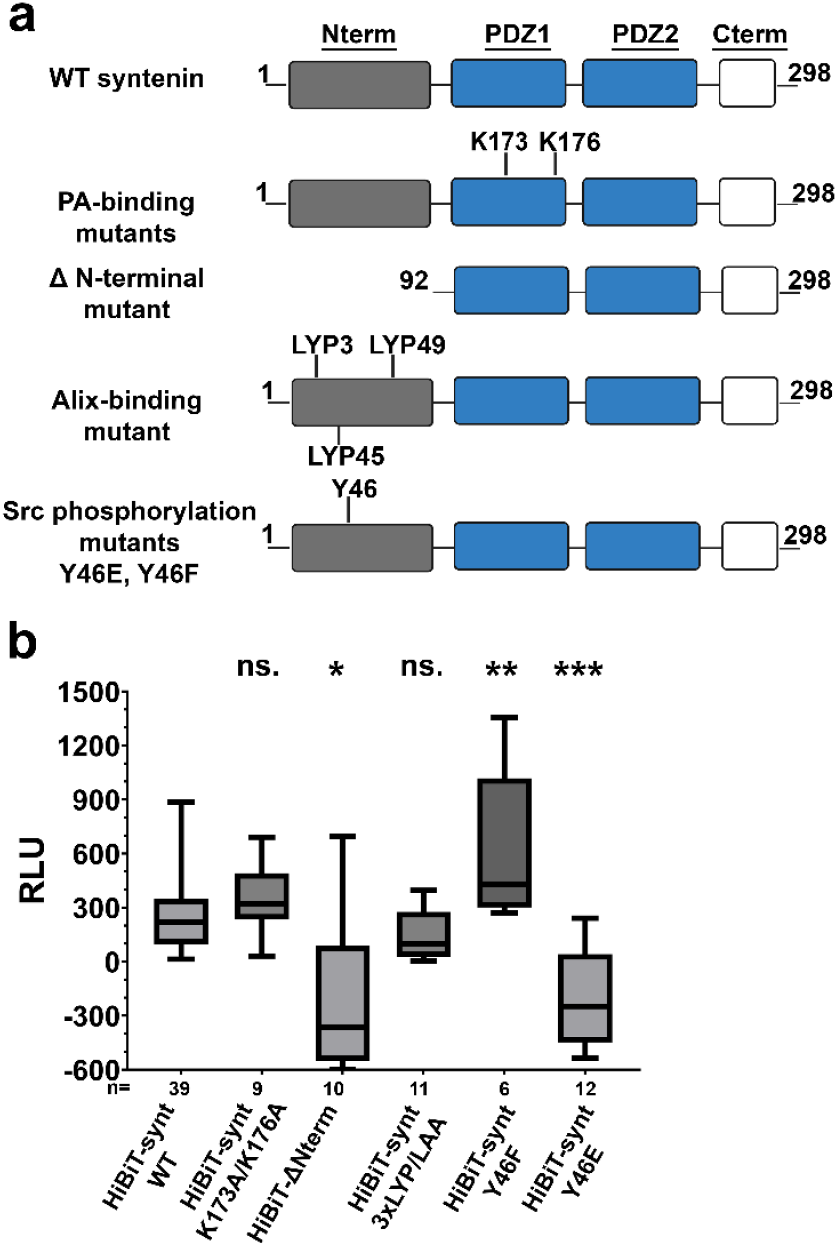
Mapping of syntenin molecular determinants involved in cytosolic delivery. **(a)** Schematic representation of syntenin protein. The N-terminal domain is in grey, PDZ domains are in blue, and C-terminal domain is in white. The wild-type form (upper scheme) and the various mutants tested in this study are represented. **(b)** Bars and whiskers graphs showing the median luminescence signals with minimal and maximal values, obtained with CCM sEVs containing HiBiT-syntenin WT or different HiBiT-syntenin mutants as indicated. Background signals from MCF-7 LgBiT cells treated with Mock CCM sEVs were subtracted. Three to 15 different preparations of syntenin WT or mutant CCM sEVs were used (Supplementary Fig. 6b) to collect 6-39 independent measurements, as indicated. One-way ANOVA with Dunnet’s correction for multiple comparisons was applied, comparing WT syntenin cytosolic signal with that of the mutants. Each CCM sEV preparation (see Supplementary Fig. 6b) served for 1-3 independent measurements of cytosolic delivery. Related to **Supplementary Fig. 6**.

Next, we established experimental conditions allowing to quantify the amount of HiBiT in sEVs by luminescence. We observed that, in the presence of 1.66 μM Furimazine (membrane permeant substrate for NL), 0.05% Triton X-100 detergent, and 2.5 μg rLgBiT (recombinant form used to saturate HiBiT), luminescence signals were directly proportional with the reagent dilution (Supplementary Fig. 4b, c). To determine optimal reagent concentrations (Fig. 2c), we tested different dilutions of CCM sEVs and decided to use a 1/20 dilution (within the linear range), to measure the total sEV dose in subsequent experiments.

CCM sEVs were then incubated in the presence (0.05%) or absence of Triton X-100 (Fig. 2d). Luminescence was detected only when both detergent and rLgBiT were added (Fig. 2e). These results indicate the HiBiT-syntenin is completely embedded within sEVs. Similar observations were obtained for HiBiT-CD63, consistent with the cytosolic orientation of its N-terminus. Full-length NL fused to syntenin served as a positive control.

Together, these data establish that HiBiT-syntenin is strictly embedded within CCM sEVs and can be robustly quantified by luminescence following sEV permeabilization and NL complementation.

### Establishing conditions for measuring internalized sEVs

To distinguish between ‘captured’ (retained, but non-internalized) and internalized sEVs, we treated the recipient cells with trypsin. As we and others documented that heparan sulfate proteoglycans (HSPGs) are major receptors for sEV uptake^6,21^, and as HSPGs are highly trypsin sensitive, we expected that trypsin treatment would remove cell-surface captured sEVs. Recipient MCF-7 cells were therefore first incubated with NL-syntenin CCM sEVs, washed to remove unbound vesicles, and then treated with trypsin and washed again. When NL-syntenin CCM sEVs (30p/r) were incubated with MCF-7 wild-type (WT) cells, luminescence signals measured after the washing of the cells in the absence of trypsinization continued to increase over time, without reaching equilibrium. In contrast, when cells were treated with trypsin and washed, luminescence signals were reaching a plateau after 60 minutes and remained stable for up to 4 hours (Fig. 3a).

These results indicate that a substantial fraction of the signal measured without trypsinization originates from ‘captured’ sEVs, bound to or trapped between cells.

Next, to address cytosolic delivery, we generated MCF-7 cells stably expressing cytosolic LgBiT (Supplementary Fig. 5a). We first tested whether NL complementation could be detected upon transfection with expression vectors encoding HiBiT-syntenin or HiBiT-CD63. Empty vector (Mock) was used as a negative control and NL-syntenin was used as a technical control (Supplementary Fig. 5b). Although Mock-transfected cells displayed high background luminescence, signals were more than two orders of magnitude higher in cells transfected with expression vectors encoding HiBiT-syntenin or HiBiT-CD63, confirming that NL complementation can occur and can be measured.

We next set up experimental conditions to measure the total internalized (not necessarily release in the cytosol) syntenin sEV fraction. We found that Triton X-100 concentrations between 0.5–2%, in the presence of 2.5 μg rLgBiT, provided stable and reproducible luminescence signals (Supplementary Fig. 5c, d).

Taken together, these set-up experiments indicate that it is possible to measure reconstituted NL luminescence in MCF-7 LgBiT cells, including in the presence of detergent.

### Internalization and cytosolic delivery of sEV HiBiT-syntenin in MCF-7 cells expressing LgBiT

Having defined conditions to measure the internalized syntenin sEV fractions, we incubated recipient MCF-7 LgBiT cells with HiBiT-syntenin CCM sEVs for 4 hours (30p/r) and compared three distinct experimental parameters: (i) the total administered HiBiT sEV dose, (ii) the internalized fraction (measured after trypsinization and detergent permeabilization), and (iii) the cytosolic fraction (measured after trypsinization in the absence of detergent) (Fig. 3b).

Luminescence signals are shown in Fig. 3c (raw data in Supplementary Table 2). For HiBiT-syntenin, quantitative analysis revealed that only ∼0.03% of the administered CCM sEV dose was internalized by MCF-7 LgBiT cells within 4 hours, whereas cytosolic delivery represented 84.7 +/-36.3% of the internalized sEV fraction. Although variability between experiments was observed, cytosolic delivery consistently represented a large fraction of the internalized pool across biological replicates. For NL-syntenin CCM sEVs (technical control), signal corresponding to ∼0.01% of the total dose was detected following detergent permeabilization of MCF-7 LgBiT cells, while 70.8 +/-8.5% of the internalization signal was detected in the absence of detergent (Fig. 3c). For HiBiT-CD63 (used as positive control), the internalization was ∼0.02% compared to the full dose and cytosolic delivery was 31.9 +/-12.4% of the internalized fraction (Supplementary Fig. 5e).

Taken together, these results show that solely a small fraction of the administered sEV dose is internalized, while a large fraction of the internalized pool undergoes cytosolic delivery.

### Syntenin molecular determinants supporting cytosolic delivery

We then investigated molecular determinants within syntenin that support cytosolic delivery. Fortunately, mutants were found in equivalent amounts in CCM sEVs, as sEV HiBiT-syntenin total dose measurements were similar for WT and mutant constructs (Supplementary Fig. 6b, raw data in Supplementary Table 2).

First, we examined whether binding of syntenin to the fusogenic lipid phosphatidic acid (PA), via its PDZ1^8^, contributes to cytosolic delivery. A syntenin mutant containing two lysine-to-alanine substitutions within the putative PA-binding site of PDZ1 was generated (Fig. 4a, Supplementary Fig. 6a). These mutations did not significantly alter cytosolic delivery efficiency in recipient cells compared to WT syntenin (Fig. 4b). We next investigated the role of the N-terminal domain, as this domain interacts with Alix, and a report has implicated Alix in back-fusion of intraluminal vesicles with the limiting endosomal membrane^22^. We first prepared a syntenin mutant with deletion of this domain (ΔNterm). Cytosolic delivery of this mutant was below the background signals obtained with Mock CCM sEVs, suggesting that deletion of the N-terminal domain completely abolishes cytosolic delivery.

Next, we tested an Alix-binding-deficient syntenin mutant in which the three N-terminal LYP motifs were replaced by LAA^5^. Unexpectedly, this mutation did not affect cytosolic delivery efficiency compared to WT syntenin (Fig. 4b). Finally, because the N-terminal domain also contains the Src phosphorylation site tyrosine-46^11^, we also tested the cytosolic delivery of Src phosphorylation-deficient (Y46F) or phosphorylation-mimetic (Y46E) syntenin mutants. Surprisingly, cytosolic delivery of the Y46F mutant was significantly increased compared to WT syntenin, whereas cytosolic delivery of the Y46E mutant was completely abolished (Fig. 4b).

Taken together, these results indicate that the N-terminal domain of syntenin is essential for cytosolic delivery and that the phosphorylation status of tyrosine-46 functions as a molecular switch, with phosphorylation inhibiting and dephosphorylation promoting cytosolic delivery.

## Discussion

In this study, we demonstrate that sEV-embedded syntenin undergoes delivery to the cytosol of recipient cells. Using a GFP complementation assay, we observed cytosolic delivery of sEV-embedded syntenin in ∼9% of recipient cells within 4 hours (Fig. 1d) upon incubation with relatively low amounts of GFP11-syntenin CCM sEVs (secretome of 30 producing cells per recipient cell (30p/r). This relatively high fraction of GFP-positive cells is striking given the generally low efficiencies reported for sEV-mediated cargo transfer. While one could argue that high efficiency of GFP reconstitution might result from transfer of DNA encoding GFP11-syntenin^23^, the short incubation time, together with the requirement for cytosolic protein complementation, makes de novo transcription, translation, and subsequent GFP reconstitution highly unlikely. In other words, we are confident that GFP reconstitution can be solely explained by the transfer of GFP11-syntenin protein from donor cells to the cytosol of recipient cells expressing GFP1-10.

Our GFP reconstitution assays also highlight that the method of sEV preparation may influence syntenin delivery. Noteworthy, we observed that recipient cells treated with GFP11-syntenin CCM sEVs displayed nuclear and cytosolic dotty GFP patterns. These cytosolic dots may be explained by efficient local GFP reconstitution driven by membrane-associated clustering of GFP11-syntenin, consistent with the known enrichment of syntenin in microsomal fractions^2^ and its interaction with various lipids^8,24,25^. Accordingly, GFP reconstitution mediated by cytosolic GFP11-syntenin molecules that are not membrane-associated may remain below the detection threshold of this microscopy assay. Interestingly, both dotty and nuclear distributions are reminiscent of endogenous syntenin distribution^2^.

To complement this spatially resolved but poorly quantitative approach with a more sensitive and flexible approach, we used Nanoluciferase (NL) and a NL-complementation assay^26^. Using this assay, we first established that HiBiT-syntenin is strictly embedded within sEVs (Fig. 2d), thereby excluding non-vesicular delivery of syntenin. Using NL-syntenin sEVs we show that capture exceeds internalization by nearly one order of magnitude within the first hours (Fig. 3a), and fails to reach equilibrium over time. Heparan sulfate proteoglycans are thought to play a prominent role in the capture and internalization of sEVs^7,21^. Because recipient cells were freshly plated prior to sEV addition, it is plausible that heparan sulfate proteoglycan abundance or accessibility at the cell surface increases over time, which may contribute to the progressive increase of the captured signal. As heparan sulfate proteoglycans are highly protease-sensitive^27^, trypsin digestion is expected to remove (most of) the capture-associated signal. The remaining signal (here considered as internalized fraction), in contrast to capture, rapidly reaches a plateau value and remains stable over time (Fig. 3a). Internalization reaching rapid equilibrium implies that the internalized (NL) cargo is metabolically unstable (i.e. is degraded and/or is again leaving cells). Thus, (at least here) capture dominates early sEV-cell interactions and can mask the comparatively limited internalization process. This separation between capture and internalization is critical for interpreting sEV delivery measurements. Notably, while we previously showed that total ‘uptake’ (capture plus internalization combined) can be saturated when high sEV doses are applied^14^, whether increasing sEV dose also enhances internalization remains an open question.

After clarifying the importance of separating sEV capture from internalization, we incubated CCM sEVs with recipient MCF-7 LgBiT cells and found that (at equilibrium) the internalized HiBiT-syntenin corresponds to only a very small fraction (∼0.03%) of the administered HiBiT-syntenin sEV dose. Yet relative to this internalized fraction, cytosolic delivery of this syntenin was extremely effective (85%) (Fig. 3c).

These observations were unexpected in light of the studies stating that cytosolic delivery is negligible in the absence of viral fusogenic proteins, whereas the internalization is efficient^20,28–30^. Of note, these conclusions are often based on microscopy-based readouts that do not measure the total administered dose^30^, lack correction for background luminescence^29^, or do not distinguish between capture and internalization^31^.

In contrast, our findings are consistent with those of Bonsergent et al., who similarly concluded that internalization, rather than cytosolic delivery, represents the limiting step of sEV cargo delivery^32^. That study did not use split reconstitution assays but used cell fractionation and an HSP70 NL fusion protein. Of note, syntenin was shown to govern HSP70 loading into sEVs^5^. As such HSP70 and syntenin sEVs may be very similar^33^, and the efficient cytosolic delivery observed for HSP70 may therefore reflect intrinsic properties of the syntenin-dependent sEV subtype, which can represent up to 50% of sEVs^5,11^. Together, these data clearly indicate that cytosolic delivery does not necessarily require viral fusogenic proteins.

We also started to address the molecular mechanisms supporting sEV syntenin cytosolic delivery in sEV recipient cells. Given the established role of phosphatidic acid (PA) binding in syntenin-dependent sEV biogenesis via the PDZ1 domain, and the suggested role of phosphatidic acid in membrane fusion events^34–36^, we investigated whether syntenin-PA interaction may contribute to cytosolic delivery but found no evidence supporting this hypothesis (Fig. 4b). In contrast, deletion of the N-terminal domain abolished syntenin cytosolic delivery, suggesting that this domain is required either to generate cytosolic delivery-competent sEVs or to support cytosolic delivery following internalization. Of note, syntenin lacking its N-terminal domain was also shown to promote sEV biogenesis^37^, but based on our data, these sEVs would most probably be cytosolic delivery-incompetent. The N-terminal domain is responsible for Alix binding, which has been proposed to be involved in intraluminal back-fusion and thereby may convey cytosolic delivery^22^. Nevertheless, when syntenin is mutated at its Alix binding sites, it still efficiently undergoes cytosolic delivery. Knock-out of endogenous syntenin may be required to completely exclude a role for Alix as both mutant and endogenous syntenin may belong to the same membrane domains. Interestingly, the N-terminal domain also contains tyrosine-46, which we previously identified as a Src phosphorylation site^11^. A phosphorylation-mimetic mutation (Y46E) abolishes cytosolic delivery, whereas a phosphorylation-deficient mutation (Y46F) promotes cytosolic delivery. These data indicate that tyrosine-46 functions as a molecular switch for cytosolic delivery and suggest that Src may dampen the ‘fusogenic’ properties of sEVs toward recipient cells. Considering that Src promotes pro-migratory activities of sEVs^11^, a corollary of these observations is that the pro-migratory activity of sEVs may be unrelated to their ‘fusogenic’ activity, and thus to the release of specific molecules present inside sEVs. By extension, the pro-migratory activity of sEVs may instead rely on contact-dependent signaling and/or metabolic effects after lysosomal degradation. These exciting considerations require further investigations.

We also assessed cytosolic delivery of CD63. Although CD63 and syntenin directly interact^38^ and may thus be present on the same sEV subtype, CD63, compared to syntenin, showed lower cytosolic delivery as measured by NL complementation (Supplementary Fig. 5e). This might be due to HiBiT tag cleavage (Fig. 2b), which can render complementation inefficient even when these sEVs reach the cytosol. Such cleavage, together with indirect cytosolic delivery measurements (e.g. using Cre activity as a cytosolic delivery readout) or overestimation of internalization, may explain the inconsistent CD63 cytosolic delivery efficiencies^17,18,28,30,39^. Alternatively, for reasons that remain unclear, CD63 release from sEVs may be inefficient, consistent with the need for CD63-cleavable constructs reported by others^30,40,41^.

Independently of these conceptual considerations, our study also establishes a quantitative assay to measure sEV-mediated internal cargo delivery. However, our experience demonstrates that several technical aspects can be laborious and should be carefully considered before implementation. In particular, as detergent is required to quantify both the total dose and internalized fraction of sEV HiBiT-syntenin, experiments should be carefully set-up. Indeed, when Triton X-100 is used, luminescence increased at low detergent concentrations before returning to baseline at higher levels (Supplementary Fig. 3d), highlighting the need to define appropriate detergent ranges. Endogenous LgBiT levels in recipient cells were insufficient to support NL reconstitution under these conditions (Supplementary Fig. 3e), necessitating recombinant LgBiT supplementation. Luminescence derived from reagents also creates background signal, underscoring the need for stringent controls. Together, these considerations emphasize that although NL complementation provides high sensitivity, careful optimization is essential for its reliable application to sEV delivery studies.

In conclusion, we identify a novel function of syntenin as a mediator of sEV cytosolic delivery and establish a quantitative assay for assessing this delivery. Together, our findings illustrate that the method of sEV preparation affects sEV internal cargo delivery and, by extension, likely sEV function. They indicate that sEV-syntenin capture is slow in reaching equilibrium and (although specific) may thus be more complex than simple cell surface-binding; that internalization is rapid in reaching equilibrium, but minimal when considering the administered dose; and that syntenin supports efficient cytosolic delivery through a tyrosine located in its N-terminal domain, which functions as a switch. Clearly, syntenin supports cytosolic delivery without requiring fusion to viral proteins, highlighting its potential as a biologically relevant scaffold for sEV-mediated cytosolic cargo delivery. Our study also supports the notion that enhancing sEV internalization may help increasing the overall sEV delivery. That issue may thus be the first priority from an EV-therapeutic based perspective.

## Data availability statement

The data generated in this study are available from the corresponding author upon request.

## Acknowledgements

The authors thank Antonio-Egea Jimenez for mapping of the syntenin putative PA-binding domain. We also thank Clotilde Thery for useful discussions regarding our work. This research project was funded by Kom op tegen Kanker (Stand up to Cancer), the Flemish cancer society (projectID: 12367) and the internal funds of the KU Leuven (C14/20/105). L.H. is a PhD Fellow of the Research Foundation – Flanders (FWO, 1SE2422N). Schemes were created using BioRender.com.

## Author contribution

P.Z., G.D. and L.H. conceived and designed the study and wrote the manuscript; L.H., E.J and S.M. performed experiments and analyzed the data. G.D., C.T, A.J, P.Z., and L.H. provided critical suggestions and revised the manuscript. All authors read and approved the final manuscript.

