## Supplementary figures and legends for "Measuring capture, internalization and cytosolic delivery of extracellular vesicle-embedded syntenin"

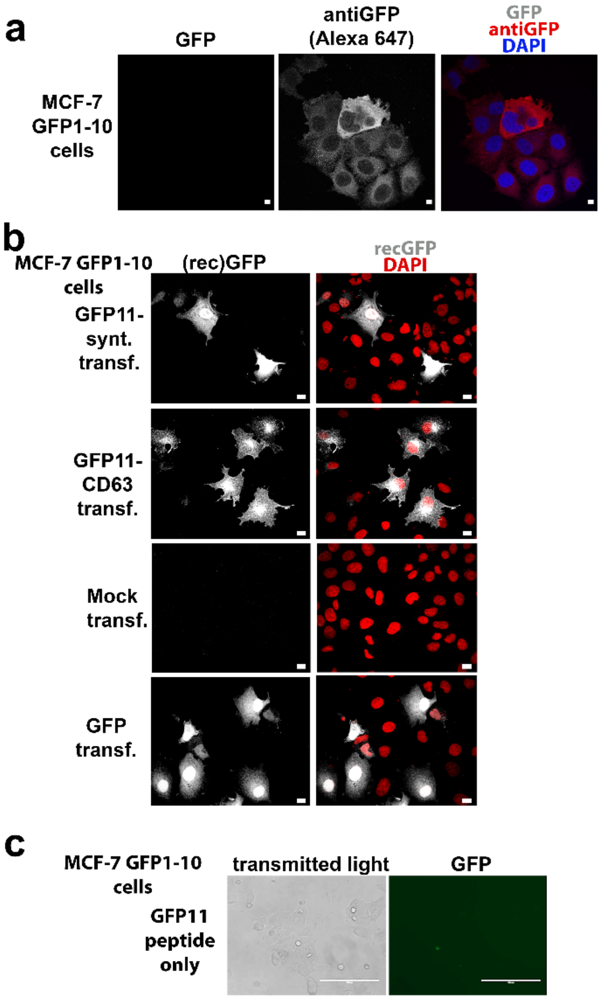


Supplementary Figure 1

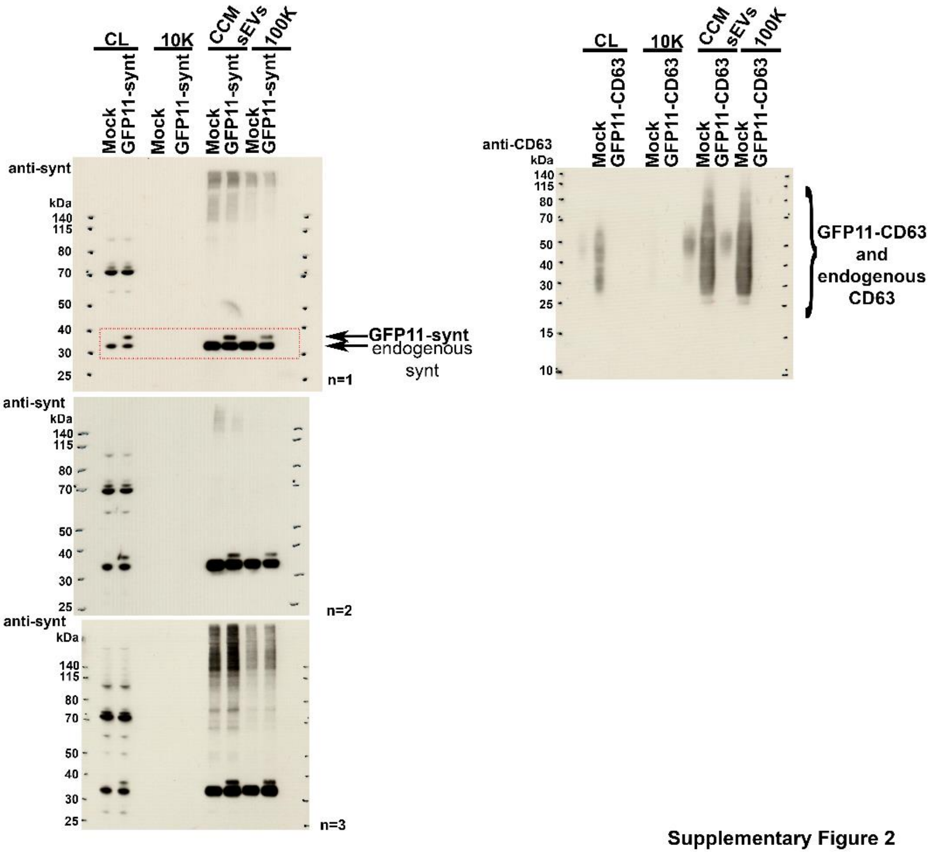


Supplementary Figure 2

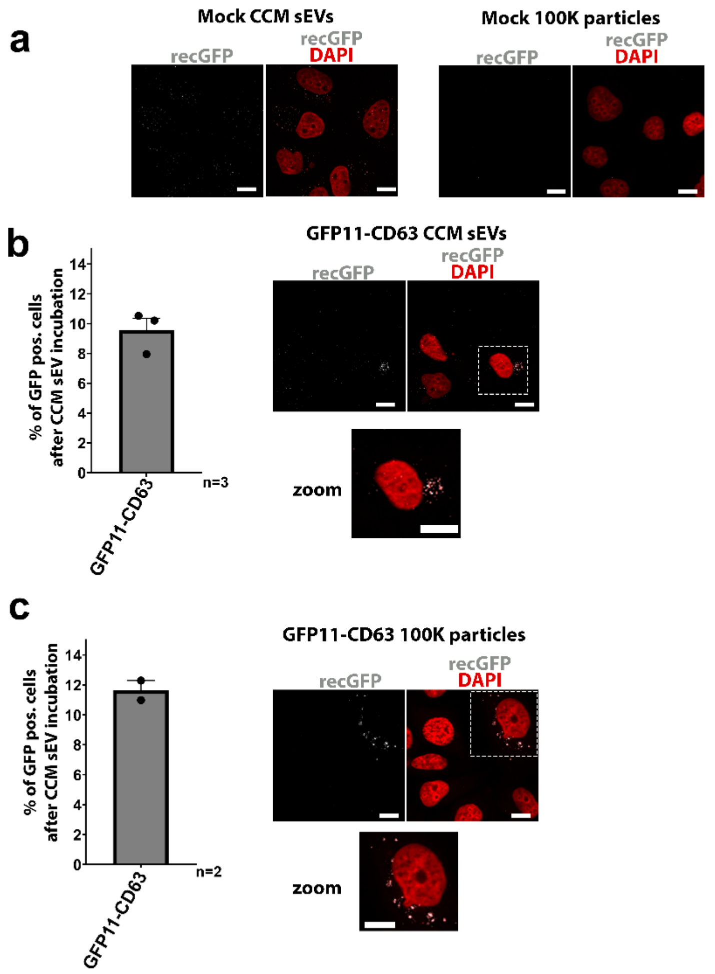


Supplementary Figure 3

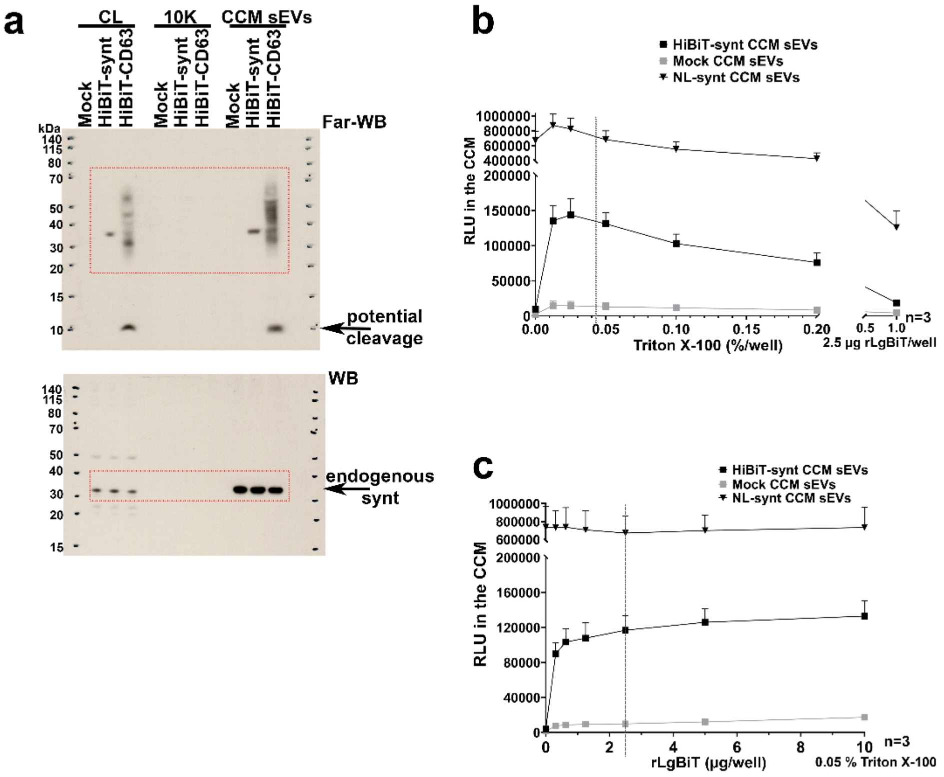




Supplementary Figure 4

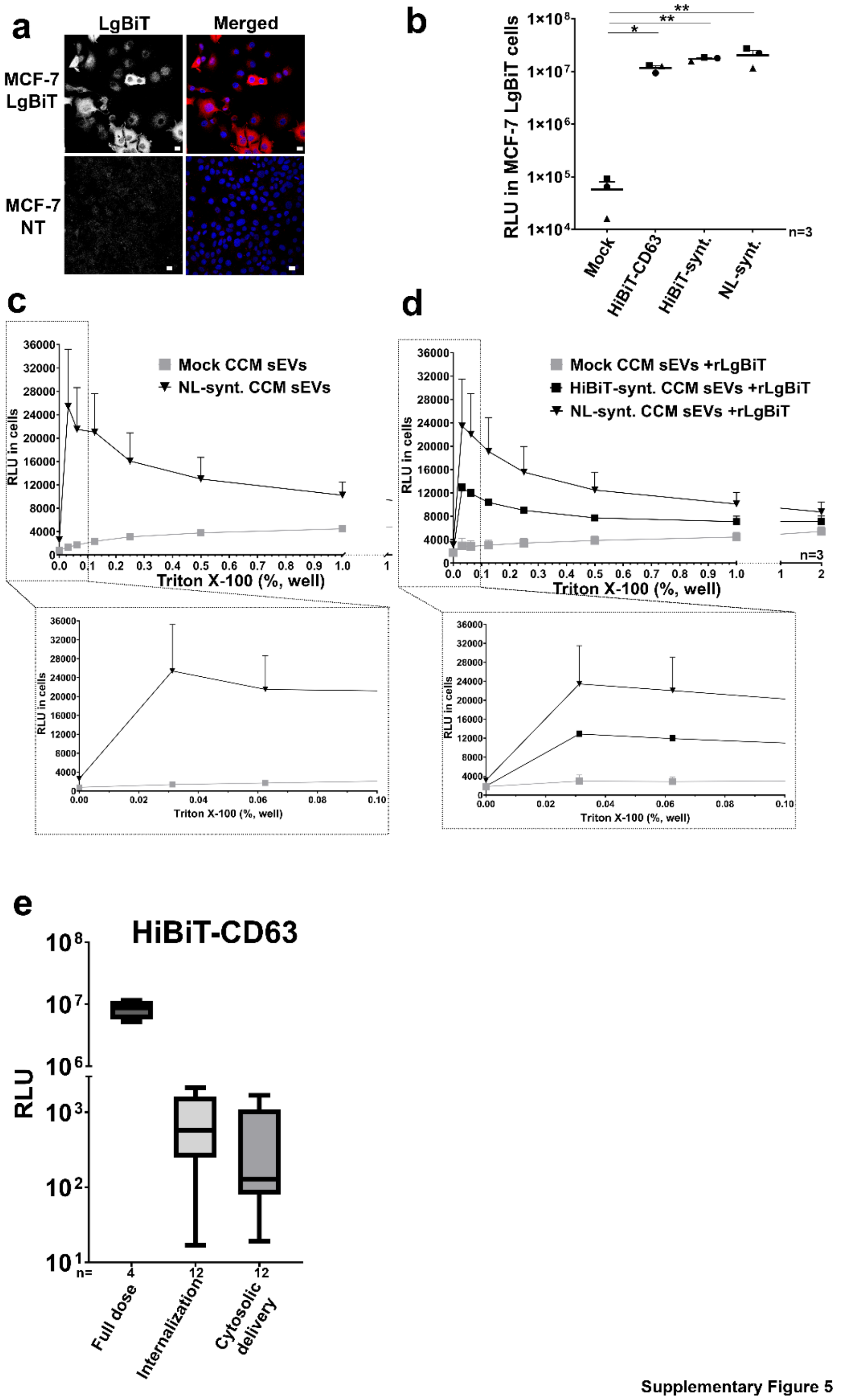


Supplementary Figure 5

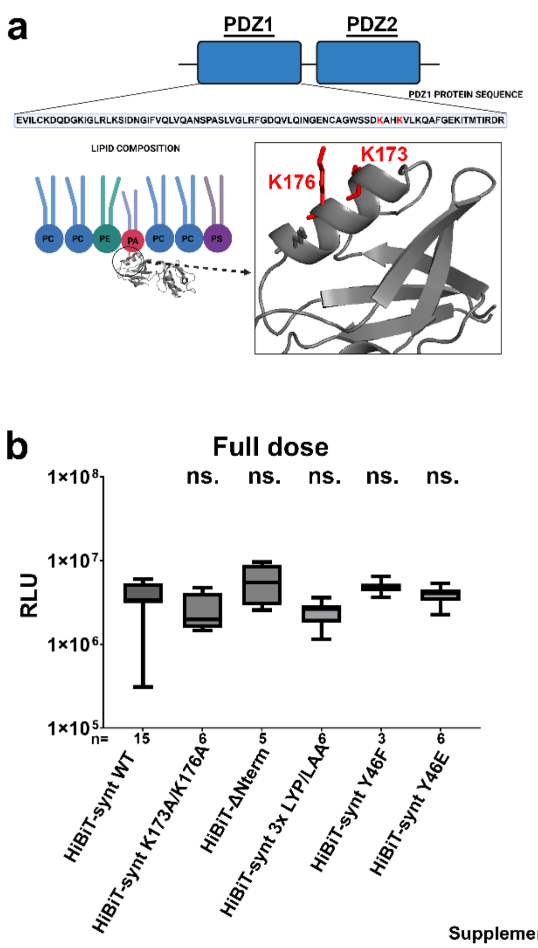


**Supplementary Figure 6**

### Supplementary figure legends

**Supplementary Figure 1. Characterization of MCF-7 cells stably expressing GFP1-10 and validation of the GFP complementation assay. (a)** MCF-7 cells stably expressing GFP1-10 were characterized by confocal microscopy using anti-GFP antibodies and Alexa-647 secondary antibodies. As expected, no fluorescence was observed in the green channel (GFP). Note that all MCF-7 cells are positive for GFP1-10 expression. Scale bar is 10 µm. **(b)** Confocal micrographs of MCF-7 GFP1-10 cells showing reconstituted GFP fluorescence after transfection with an expression vector for GFP11-syntenin (upper part), GFP11-CD63 (upper middle part), Mock transfected (lower middle part) or full-length GFP used as positive control (lower part). Scale bar is 10 µm. **(c)** Wide field micrographs (transmitted light left, fluorescence right) of MCF-7 GFP1-10 cells transfected with an expression vector for GFP11 peptide. Note the complete lack of GFP fluorescence consistent with the instability of the GFP11 peptide when taken in isolation. Related to **Fig. 1.**

**Supplementary Figure 2. Uncropped and additional Western blots showing the expression of GFP11-fusion constructs in cell lysates and secretomes.** Western blots of HEK293 cell lysates (CL) and their secretomes fractioned by centrifugation to remove lEVs (10K) and followed by concentration (CCM sEVs) or differential ultracentrifugation (100K), as indicated. HEK293 were transfected with GFP11-syntenin (left) or GFP11-CD63 (right) and probed with anti-syntenin or anti-CD63 antibodies, respectively, recognizing both the endogenous and the GFP11-tagged forms. CL corresponds to 20,000 cells and secretomes correspond to 2.4 million cells. Related to **Fig. 1.**

**Supplementary Figure 3. GFP complementation assay showing the MCF-7 GFP1-10 cells after incubation with Mock or GFP11-CD63 CCM sEVs and 100K particles. (a)** Confocal micrographs showing the lack of GFP fluorescence (grey) in MCF-7 GFP1-10 treated with Mock CCM sEVs (left) or 100K particles (right). Nuclei were stained with DAPI (red). Scale bar is 10 µm. **(b, c)** Left, bar graphs showing the average values + SEM of GFP positive MCF-7 GFP1-10 cells after the incubation with GFP11-CD63 CCM sEVs **(b)** or 100K particles **(c)**. Each point corresponds to an independent biological repeat from at least 40 nuclei counted at random. Right, confocal micrographs showing the localization of the GFP fluorescence (grey) in MCF-7 GFP1-10 cells treated with GFP11-CD63 CCM sEVs **(b)** or 100K particles **(c)**. Nuclei were stained with DAPI (red). Inserts correspond to higher magnification of specific areas, as indicated. Note dotty cytoplasmic signals of the reconstituted GFP. Scale bar is 10 µm. Related to **Fig. 1.**

**Supplementary Figure 4. Uncropped Western blots and determination of the experimental conditions to measure the dose of HiBiT-fusion constructs encapsulated in sEVs. (a)** Uncropped western blots related to Fig. 2B. Red squares indicate cropped areas. **(b, c)** Graphs showing average luminescence signals (RLU, Y-axis) + SEM obtained with HiBiT-syntenin, Mock and NL-synt CCM sEVs incubated in the presence of 2.5 µg of rLgBiT per well and various concentration of Triton X-100 **(b)** or various amounts of rLgBiT per well in the presence of 0.05% Triton X-100 **(c)**. Dashed lines indicate selected experimental conditions for further total dose measurements. Namely, 0.05% Triton X-100 and 2.5 µg of rLgBiT per well. Note the minimal background obtained for Mock CCM sEVs. The n-number indicates the number of independent biological repeats. Related to **Fig. 2.**

**Supplementary Figure 5. Characterization of MCF-7 cells stably expressing LgBiT, validation of NL complementation assays in these cells, determination of the experimental conditions to measure reconstituted NL activity and measurements (dose, internalization, cytosolic delivery) with HiBiT-CD63 CCM sEVs. (a)** MCF-7 cells stably expressing LgBiT were characterized by confocal microscopy using anti-LgBiT antibodies and Alexa-647 secondary antibodies. As expected, no fluorescence was observed in non-transfected (NT) cells. Cells were stained with DAPI (blue) to identify nuclei. LgBiT is red in merge. Note that all MCF-7 cells are positive for LgBiT expression. Scale bar is 30 µm. **(b)** Dot plot showing luminescence signals (RLU, Y-axis) in MCF-7 LgBiT cells transfected with HiBiT-CD63, HiBiT-syntenin, NL-syntenin (used as positive control), or empty vector (Mock, used as negative control), as indicated. The n-number indicates the number of independent biological repeats. Each point corresponds to an average of at least two technical repeats; lines correspond to mean + SEM. **(c, d)** Graphs showing mean luminescence signals (RLU, Y-axis) + SEM observed with MCF-7 LgBiT cells that were incubated (4 hours, 30p/r) with Mock, NL-syntenin **(c-d)** or HiBiT-syntenin **(d)** CCM sEVs. After trypsinization, cells were incubated with varying concentrations of Triton X-100 in the absence **(c)** or presence **(d)** of rLgBiT. Background signals obtained upon incubation with Mock CCM sEVs are shown in grey. Inserts allow to better distinguish signal variations obtained with low Triton X-100 concentrations (0-0.1%). The n-number indicates the number of independent biological repeats. **(e)** Bars and whiskers graphs showing the median luminescence signals with minimal and maximal values, obtained with CCM sEVs containing HiBiT-CD63. Background signals from MCF-7 LgBiT cells treated with Mock CCM sEVs were subtracted. Each CCM sEV preparation served for 3 independent measurements of cytosolic delivery. Related to **Fig. 3.**

**Supplementary Figure 6. Representation of the putative phosphatidic acid binding site in syntenin PDZ1 domain and measurement of the full dose of HiBiT-syntenin WT and mutants. (a)** Modeling of the putative syntenin-phosphatidic acid binding sites. Residues 171-173 are predicted to interact with phosphatidic acid. **(b)** Bars and whiskers graphs showing the median luminescence signals with minimal and maximal values of the administered CCM sEV dose, obtained with CCM sEVs containing HiBiT-syntenin WT or mutants, as indicated. Background signals from Mock CCM sEVs were subtracted. Three to fifteen different preparations of syntenin mutant

CCM sEVs were used, as indicated. One-way ANOVA with Dunnet’s correction for multiple comparisons was applied comparing WT syntenin signal with the signals of the mutants. Related to **Fig. 4.**
